# Human Rhinovirus 16 infection of HeLa-Ohio and HEp-2 cells modifies micro RNA expression across the viral life cycle

**DOI:** 10.1101/2025.07.01.660546

**Authors:** Emily Smith, Ahmed Alrefaey, Ibemusu Otele, Aref Kyyaly, Jamil Jubrail

## Abstract

Human rhinovirus (RV) is the most frequent cause of the common cold, as well as severe exacerbations of chronic obstructive pulmonary disease (COPD) and asthma. Currently, there are no effective and accurate diagnostic tools or antiviral therapies. MicroRNAs (miRNAs) are small, non-coding sections of RNA involved in the regulation of gene expression and have been shown to be associated with different pathologies. However, the precise role of miRNAs in RV infection is not yet well established. This study aimed to analyse the impact of RV16 on miRNA expression across the viral life-cycle to identify a small panel with altered expression. We then aimed to specifically interrogate these results using our in-house developed modelling programme to identify specific genes regulated by these miRNAs during RV infection which can then be tested functionally. Our results first identified that three miRNAs, miR-155-5p, miR-140-3p, and miR-122-5p were potential biomarkers being differentially regulated at specific time points post infection. Our modelling program then linked these miRNAs to four genes (OLFML3, STAG2, SMARCA2, CD40LG) that play important roles in regulating the hosts antiviral responses and viral progression. Together, this work identifies a potential panel of biomarkers that could, along with our previous work, form clear diagnostic markers for RV16 infection and identifies specific targets that can be functionally interrogated helping to identify new cellular targets modified by RV16 infection that could inform future therapeutic design.

## Introduction

Rhinovirus (RV) is a small, non-enveloped virus with a positive-sense, single-stranded RNA genome, belonging to the *Picornaviridae* family (Rollinger and Schmidtke 2011). RV infects most adults on average twice a year. However, for patients with inflammatory airway conditions such as asthma and COPD, the virus can drive severe lower respiratory tract infections (LRTIs) in 50% of cases with ∼25% of these leading to a secondary bacterial infection (Wilkinson *et al*. 2019).

The ability of RV to cause these infections relies on its efficient life-cycle and being able to identify markers of these different stages and new therapeutic targets is of urgent priority. RV first binds to epithelial cells and other permissive cells throughout the respiratory tract using a range of receptors, including intra-cellular adhesion molecule 1 (ICAM-1) and low-density lipoprotein receptor (LDLR), to initiate its life cycle (Rollinger and Schmidtke 2011; Kim, Jang and Jang 2021). The virus then replicates within the cytoplasm, with cell lysis or exosomal release allowing it to then spread throughout the nasopharyngeal cavity and deeper into the lower respiratory tract. During this process, host immune defences are activated leading to symptom onset and often viral control for healthy individuals (Blaas and Fuchs 2016).

Given the speed of the viral life-cycle and the impact on patients’, accurate diagnosis of RV is vital in improving patient outcomes. Commonly, RV is diagnosed in clinical settings from symptom assessment with treatment focusing on management advice until the virus is cleared. This approach can be challenging due to the overlap of RV symptoms and progression with other respiratory viruses such as respiratory syncytial virus (RSV). Furthermore, accurately identifying the causative virus present is crucial for guiding treatment choices and determining the exact cause of exacerbations (Shi *et al*. 2025). To this end, miRNAs are proving interesting as potential biomarkers.

miRNAs are single-stranded, non-coding, 22-25 nucleotide RNAs involved in regulating gene expression, showing altered expression in diseases from cancer to viral infection, making them an attractive option for novel biomarkers and targeted treatment development. In RV, the role of miRNAs is critical to viral replication and disease progression with multiple studies having assessed the contribution of miRNAs to infection progression. However, these studies have mainly focused on cellular expression of miRNAs which can be less reliable than supernatant detection. Our previous findings found that 9 miRNAs demonstrated differential expression in the supernatant at 6- and 18-hours post HRV16 infection, with miR-101-3p and miR-30b-5p being the most significant and correlated with changes in specific genes (EZH2, RARG, and PTPN13). These findings showed that miRNAs could potentially be good diagnostic biomarkers, aiding in early HRV diagnosis (Bosner *et al*. 2023).

Here we undertook a non-biased analysis of miRNA expression during RV infection by using a panel that was not skewed towards specific functions. Our results show a clear and consistent modulation of miRNA expression at key time points during RV16 infections, particularly during entry and replication steps. Analysis of these findings using our modelling program that was further developed to interrogate the findings more deeply, suggests that this correlates with clear differential gene expression at these time points that limit antiviral responses whilst promoting viral replication and spread. This suggests that along with our previous findings, this panel of 6 miRNAs could be key biomarkers to diagnose RV16 infection and as they are regulated at different time points, could be useful at identifying patients before they become symptomatic. The further enhancement of our modelling programme offers a tool to interrogate this data deeply and identify gene targets that could be functionally interrogated. Overall, by establishing accurate miRNA biomarkers, it may be possible to not only determine disease presence, but also susceptibility to exacerbations, secondary infections, and disease severity.

## Methods

### HEp-2 and HeLa Ohio cell culture

Hep-2 cells were obtained from the American Type Culture Collection (ATCC) and HeLa-Ohio cells were obtained from the European Collection of Authenticated Cell Cultures (ECACC). Hep2 cells were maintained in RPMI 1640 supplemented with 10% foetal calf serum (FCS) and 1% penicillin-streptomycin and HeLa-Ohio cells were maintained in DMEM supplemented in the same way and with 1g/L glucose. Both cell lines were maintained in T75 flasks and passaged every 3 days or when 70% confluence was reached. Passaging was done as outlined previously (Jubrail *et al*. 2018; Currie *et al*. 2016).

### Human Rhinovirus 16 infection

Human Rhinovirus 16 (RV16) (VR-283, Strain 11757) was purchased from the ATCC and produced using HeLa-Ohio cells as outlined previously (Jubrail *et al*. 2018). Briefly, HeLa-Ohio cells, at a density of 1 x 10^6^, were infected with RV16 or media at a multiplicity of infection (MOI) 1 in viral media (DMEM supplemented with 1g/L glucose and 4% FCS). Flasks were agitated for 1hr at room temperature, after which viral media was added, and the flasks were incubated at 37°C and 5% CO_2_ for up to 48 hours or until 90% cytopathic effect was observed.

### Human rhinovirus 16 production and quantification

Once 90% CPE was achieved, virus was produced by three freeze-thaw cycles at – 80°C followed by high-speed centrifugation, and filtration (Jubrail *et al*. 2018). Viral stocks were aliquoted as 1ml and stored at –80°C for future use. Viral quantification was carried out using a tissue culture infective dose 50 (TCID50) assay as outlined previously (Jubrail *et al*. 2018). Briefly, HeLa-Ohio cells were seeded in 96 well plates at a density of 50000 per well and then infected with RV16 using a ten-fold serial dilution from pure HRV16 to 10^-9^, along with two control columns incubated with just viral media. Plates were agitated at room temperature for 1h and then viral media was added, and the plates incubated at 37°C and 5% CO_2_ for 5 days until cytopathic effect was seen in approximately 50% of the wells. TCID50 was then calculated using the Spearman-Karber formula.

### RV16 infection of HeLa-Ohio and Hep2 cells

Cells were seeded into either a 6-, 24-, or 96-well plate at a density of either 1 x 10^6^ per well (6 well), 250000 per well (24 well) or 10000 per well (96 well). After 24 hours, they were infected with RV16 or control media at a MOI of 1 and treated as detailed above with miRNA extraction being performed at defined time points post-infection (2-, 6-, 15-, 18-, or 24-hours).

### microRNA extraction

The Qiagen miRNeasy Serum/Plasma advanced mini kit was used to extract miRNAs from the supernatant according to the manufacturer’s instructions. Briefly, samples were thawed and centrifuged at 8000 x g before being homogenised in buffer RLT. The homogenised lysate was then treated with buffer AL before being centrifuged as above. Buffer RPL was then added to the supernatant and after 3 minutes buffer RPP was added for a further 3 minutes, before being centrifuged at 12000 x g. Next, to the supernatant an equal amount of isopropanol was added, and the solution was centrifuged at 8000 x g. Buffer RWT was then added and the sample centrifuged at 12000 x g. Finally, RNase-free water was added to the supernatant at room temperature for 1 minute before being centrifuged at 12000 x g. miRNA concentrations were determined using NanoDrop.

### Reverse Transcription and qPCR

The reverse transcription (RT) reaction was carried out using Qiagen miRCURY LNA RT Kit and consisted of 4µL 5x miRCURY RT reaction buffer, 10µL RNase-free water, 2µL 10x miRCURY RT enzyme mix, and 4µL template RNA (5ng/µL) per 20µL reaction. The samples were briefly centrifuged at 10000 x g. The reaction was run through a thermal cycler using the conditions in table 1, and cDNA was stored at - 20°C.

**Table 1:** Conditions used for reverse transcription.

| Setting | Temperature (°C) | Duration (mins) |
| --- | --- | --- |
| Pre-heat | 4 | 2 |
| Priming | 42 | 60 |
| Denaturing | 95 | 5 |
| Storage | 4 | Up to 4 days |

qPCR was performed using either BIO-RAD iQ SYBR Green Supermix or PowerTrack SYBR Green Master Mix as outlined below:

For BIO-RAD iQ SYBR Green Supermix, the following 10µL reaction was setup: 5µL SYBR Green, 1µL Primer, 1µL DNA template, 3µL RNase-free H_2_O.

For PowerTrack SYBR Green, the following 10µL reaction was setup: 5µL SYBR Green, 0.5µL Primer, 1µL DNA template, 3.5µL RNase-free H_2_O.

After reaction setup, the program was set according to the relevant protocol outlined in the below tables (table 2-3).

**Table 2:** BIO-RAD iQ SYBR Green Supermix qPCR program.

| Setting | Temperature (°C) | Duration |
| --- | --- | --- |
| Pre-heat | 95 | 3 minutes |
| Amplification | 95 | 15 seconds |
| Amplification | 60 | 40 seconds |
| Melt | 95 – 65 - 95 | 30 second increments |

**Table 3:** PowerTrack SYBR Green Master Mix qPCR program.

| Setting | Temperature (°C) | Duration |
| --- | --- | --- |
| Pre-heat | 95 | 2 minutes |
| Amplification | 95 | 15 seconds |
| Amplification | 60 | 60 seconds |
| Melt | 90 – 65 - 90 | 30 second increments |

### Program development

This program was coded using Python and interacts with two databases. miRDB was accessed through locally stored data downloaded from https://mirdb.org/download.html (miRDB v6.0, June 2019, MirTargetV4, miRbase 22), with the MyGene API being used to convert gene accession numbers to gene names. mirDIP is a more tissue-specific database accessed through an API, with the combination of databases allowing for a broad assessment of the targets prior to an in-depth literature search. The identified top 10 gene targets were then mapped to the RV phenotype using VarElect GeneCards API, allowing for more accurate determination of these miRNAs as biomarkers. VarElect is a bioinformatics tool that can identify direct and indirect links between miRNAs and genes through gene-phenotype co-occurrence analysis and literature mining, then ranking the genes based on the strength of the relationship to the phenotype. All results from these were exported as CSV files. The IDE used for this was Visual Studio Code (Version 1.99.2) running Python 3.11.9 (tags/v3.11.9:de54cf5, Apr 2, 2024, 10:12:12) [MSC v.1938 64 bit (AMD64)]).

## Results

### HRV16 modifies miRNA expression throughout infection in HeLa-Ohio cells

First, we wanted to examine if RV16 altered the expression of key miRNAs and U6 during infection in HeLa-Ohio cells. To do this, we infected HeLa-Ohio cells with RV16 or control media at an MOI 1 for up to 24 hours and collected the supernatants. Our results showed clear differential expression of all miRNAs examined across the time points, especially at 15-hours post infection, where there was a clear significant downregulation. Analysing each miRNA separately showed an interesting pattern: hsa-miR-122-5p showed initial upregulation (∼1.5 fold), then a gradual decrease until 15-hours post-infection where there was ∼2-fold downregulation followed by a roughly restoration in expression by 24-hours (Figure 1A). For hsa-miR-155-5p, there was relatively stable expression until 15-hours, when there was a significant downregulation of approximately 4-fold, before again restoring by 24 hours (Figure 1B). For hsa-miR-140-3p there was an upregulation from the 2-hour timepoint followed by a significant decrease of around 2-fold towards the 15-hour time point with the levels stabilising by 24-hours (Figure 1C). Finally, the expression of U6 snRNA showed a similar trend to the miRNAs, with high initial expression and a gradual decrease of around 1.4-fold towards 15-hours before again stabilising by 24 hours (Figure 1D). Taken together, these results suggest HRV16 significantly alters the miRNA landscape during infection to an overall similar degree.

**Figure 1:**
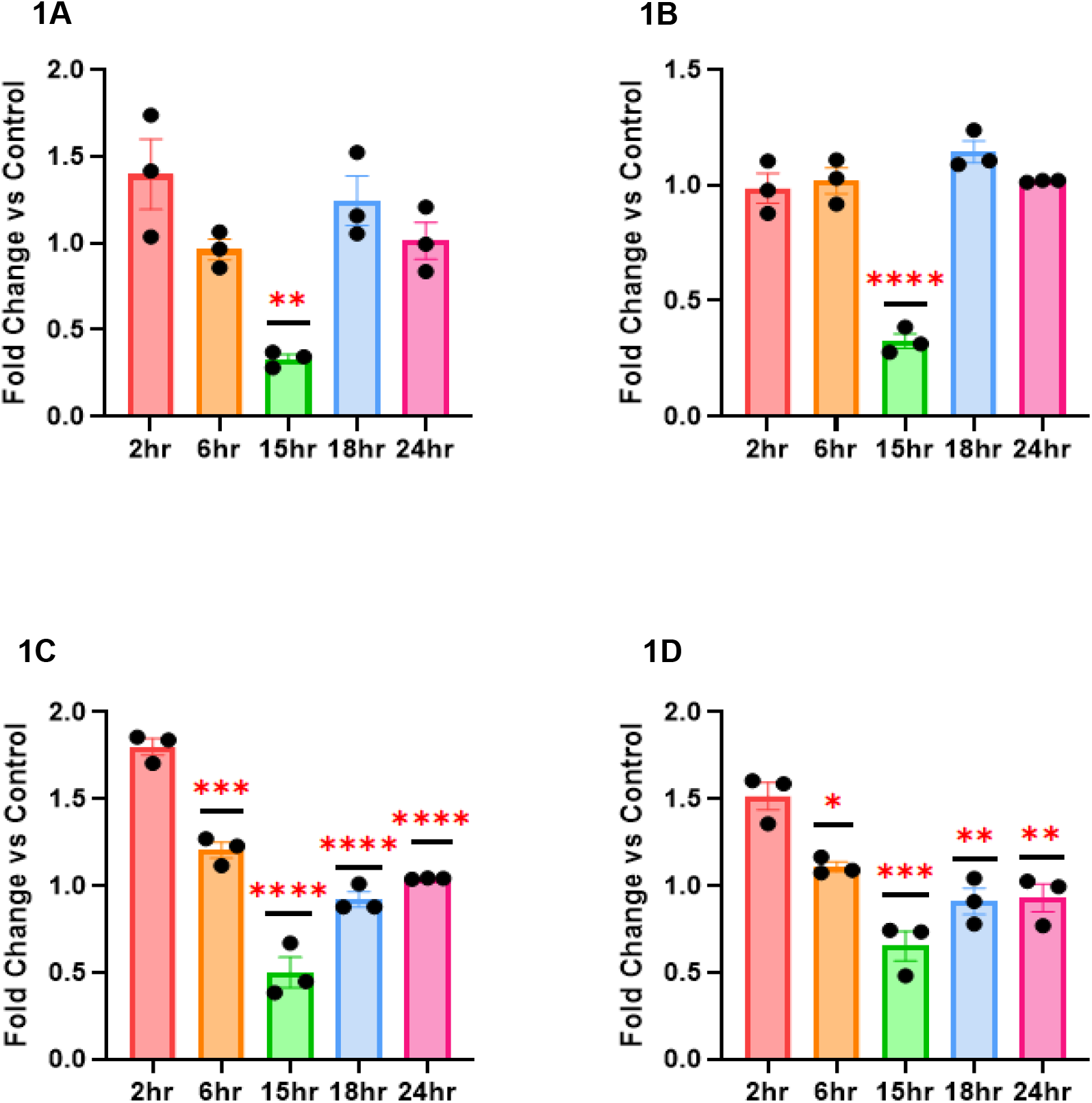
RV16 modifies miRNA expression in HeLa-Ohio cells. HeLa-Ohio cells were challenged with RV16 for different timepoints and the expression of (A) hsa-miR-122-5p, (B) hsa-miR-155-5p, (C) hsa-miR-140-3p and (D) U6 was measured by RT-qPCR. Error bars represent SEM (Standard Error of the Mean). n = 3. * P ≤ 0.05, ** P ≤ 0.01, *** P ≤ 0.001, **** P ≤ 0.0001, two-way ANOVA and post-hoc Dunnett’s multiple comparisons test vs 2 hours.

### RV16 modifies hsa-miR-155-5p and U6 expression in Hep2 cells

Based on the above results, we next assessed miRNA expression in another epithelial cell line, HEp-2. We found slight differences in the overall expression in HEp-2 cells compared to Hela-Ohio cells. For hsa-miR-122-5p, expression was relatively stable across the time points with only non-significant 1.2-fold upregulation at 2, 6 and 18 hours (Figure 2A). For hsa-miR-155-5p there was an initial downregulation at 2-hours of ∼1.8-fold, with the levels stabilising up to 24-hours post-infection when there was a significant upregulation, approximately 2.5-fold (Figure 2B). For hsa-miR-140-3p, there was no clear pattern but a trend towards a non-significant 2-fold downregulation at 15-hours post-infection followed by a second non-significant downregulation at 24 hours compared to 18 hours (Figure 2C). Finally, U6 expression showed an ∼2-fold downregulation at 2-hours post-infection before stabilising before showing a trend to a second round of downregulation at 24-hours. Taken together, these results further suggest that RV16 actively targets these miRNAs throughout its life cycle but differently depending on the cell type used.

**Figure 2:**
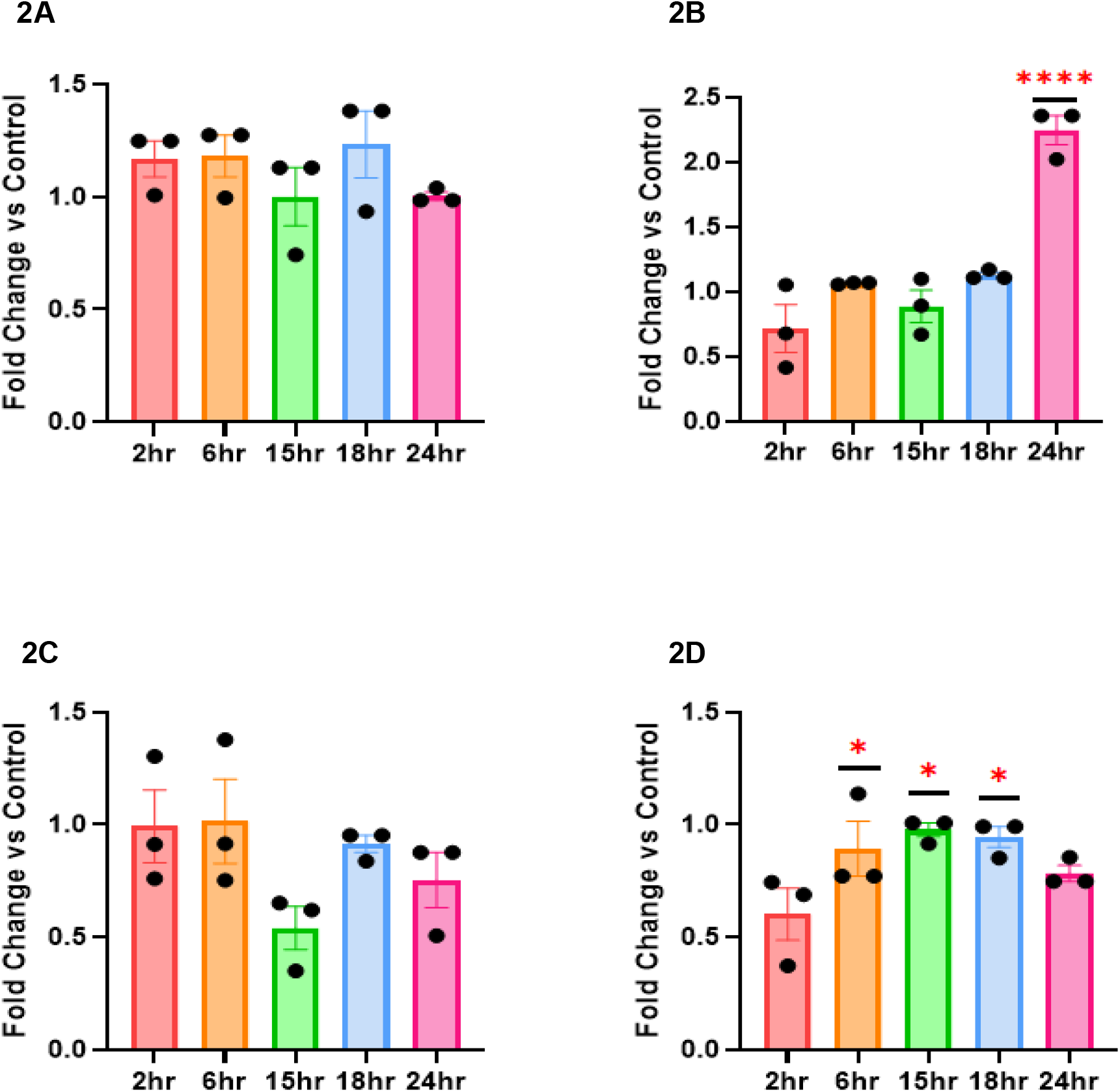
RV16 modifies miRNA expression in Hep2 cells. Hep2 cells were challenged with RV16 for different timepoints and the expression of (A) hsa-miR-122-5p, (B) hsa-miR-155-5p, (C) hsa-miR-140-3p, and (D) U6 was measured by RT-qPCR. Error bars represent SEM. n = 3. * P ≤ 0.05, **** P ≤ 0.0001, two-way ANOVA and post-hoc Dunnett’s multiple comparisons test vs 2 hours.

### RV16 targets miRNAs that regulate ten genes implicated in antiviral responses and viral replication

After these results were obtained, all three miRNAs were input into our analysis program and run through two major miRNA databases: mirDIP and miRDB v6.0. This program located the top 10 gene targets of each miRNA based on their gene scores, allowing for a broad overview of the gene targets focusing on a machine learning scoring approach (miRDB), and a more tissue-specific database (miRDIP). This allowed for a deeper view of the targets, with all resulting gene targets being input into the GeneCards VarElect API to assess relevance to the RV phenotype, via text-mining literature and gene-phenotype co-occurrence analysis, with higher scores indicating higher likelihood of significance.

The use of an API was central to the mirDIP query, allowing for streamlined data retrieval and up-to-date gene target analysis using a unidirectional search which integrates ∼30 prediction resources, some of which are tissue-specific. This API enables automated querying of this specific database. In contrast, miRDB was queried using locally stored data, downloaded from the miRDB website. After gene targets were identified (Figure 3A-C), they were input into GeneCards against the RV phenotype. This takes the [‘Gene Names’] column from the outputted CSV (comma separated values) file, and searches for the top gene results related to RV and helps to guide the overall results and potential gene targets of the miRNAs, with the results of this shown in Figure 4. The genes identified from our program suggest all three miRNAs have a significant role in RV infection through regulating specific patterns of gene expression during its life cycle.

**Figure 3:**
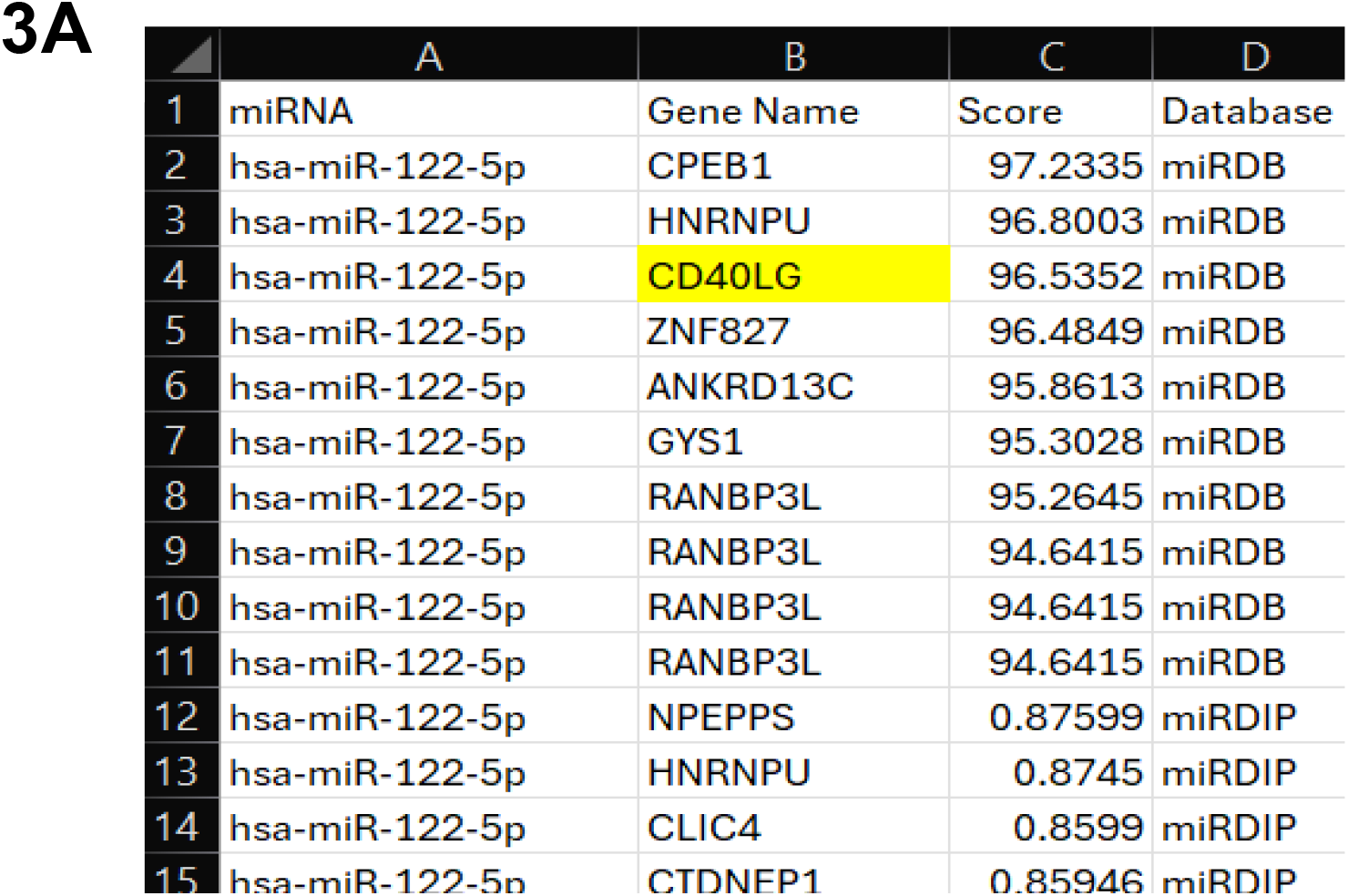
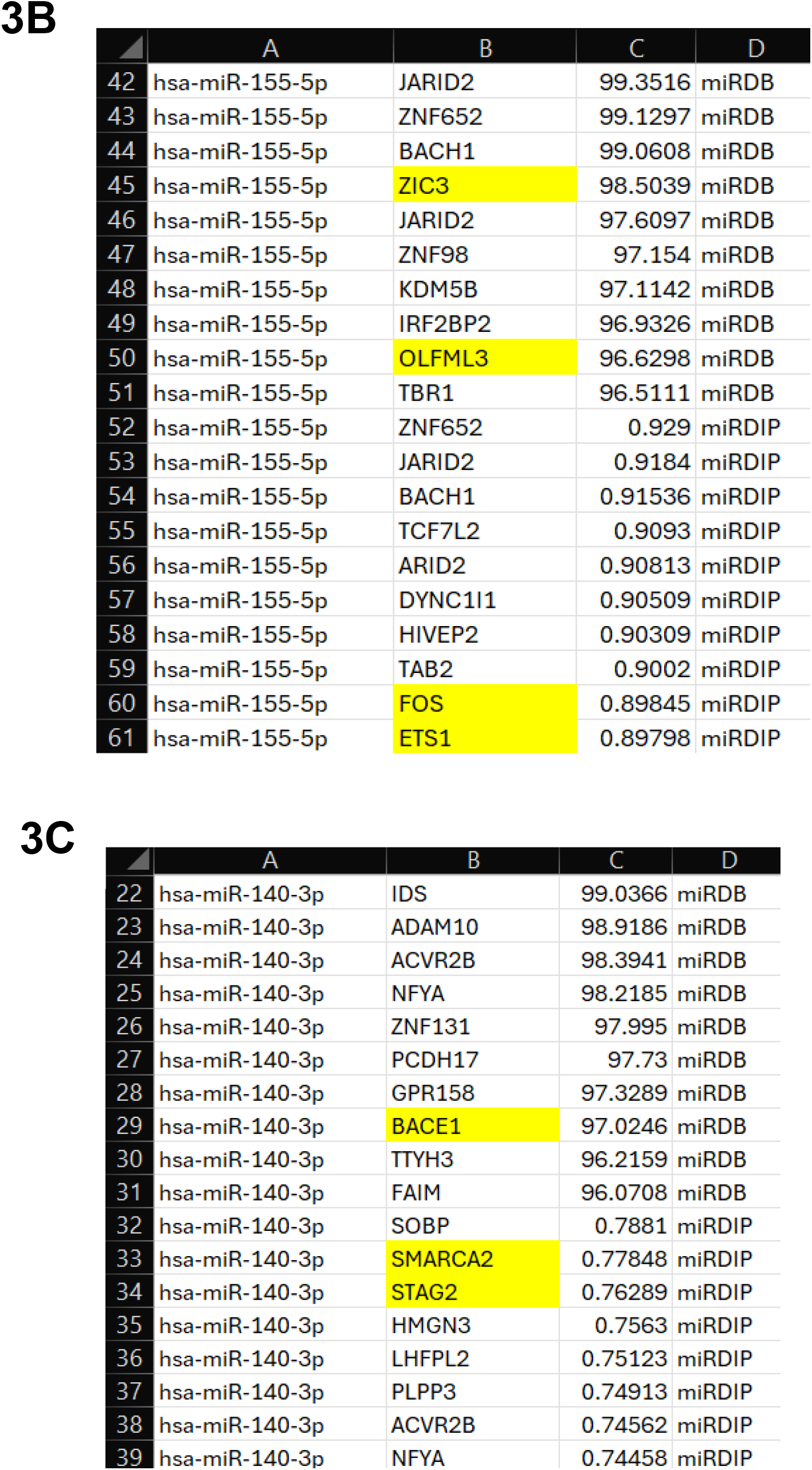
Top ten gene targets for the miRNAs of interest from miRDB and miRDIP. All three miRNAs and the desired number of gene targets (ten) was input into the program. This searched the two databases (miRDB and miRDIP) and identified the top ten targets from each, sorting them based upon TargetScore (miRDB) or IntegratedScore for the specified Score Class (miRDIP). Targets were chosen as significant if they had scores of 94+ for miRDB, and 0.70+ for miRDIP, both of which are considered as very high confidence on the respective databases. Genes identified as being linked to the RV phenotype by VarElect GeneCards are highlighted in yellow. A) Targets of hsa-miR-122-5p, with two genes (CD40LG and ALDOA) showing to be associated with this miRNA. B) Targets of hsa-miR-155-5p, with four genes (ZIC3, OLFML3, FOS, and ETS1) showing to be associated with this miRNA. C) Targets of hsa-miR-140-3p, with four genes (BACE1, SMARCA2, STAG2, and RBFOX2) showing to be associated with this miRNA.

**Figure 4:**
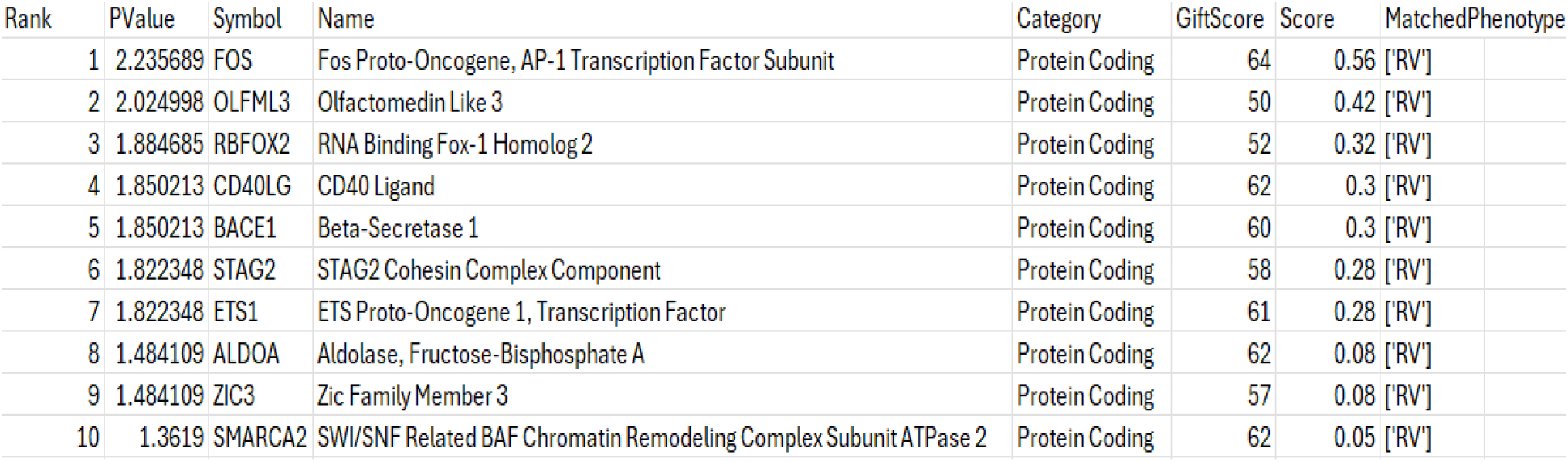
Ten genes identified as being involved in RV infection by the VarElect GeneCards API. Filtered table displaying VarElect GeneCards results for gene targets against the RV phenotype. Gene targets taken from the database search results displayed in Figure 4 (A-C). Genes are ranked by score, relating to their relevance to the RV phenotype from analysis of literature and gene-phenotype co-occurrence.

### RV16 targets antiviral responses and signalling pathways through regulation of gene expression

From the list of gene targets in Figure 4, some were disregarded due to not having enough published information linking them to the host immune response. While the API reveals a link to RV, due to the field of miRNAs still being relatively new, there are significant gaps in research in relation to their specific pathways and targets. To overcome this, searches focused on genes which had broad roles in viral infection, or where altered expression could favour/reduce HRV progression. From this, four genes (STAG2, CD40LG, SMARCA2, and OLFML3) were eventually chosen due to their high scores from the databases and their established links to the immune response and viral infection.

From our initial screen, BACE1, ZIC3, and RBFOX2 were excluded after initial literature searches revealed limited evidence of involvement in viral infections. In addition, ALDOA was removed due to insufficient evidence of their role compared to other genes regulated by these miRNAs. Finally, FOS and ETS1 were excluded as, compared to other genes regulated by hsa-miR-155-5p, there is little evidence for their direct role in viral infection. While they may be contributory to the overall response, and observed changes in miRNA expression, their role does not appear as significant as that of others such as OLFML3 (Djeddi *et al*. 2024; Aguilar-Delgadillo *et al*. 2024)

Once the four genes of interest had been identified, it became clear as to their importance within RV16 infection, and how this translates into the observed changes in miRNA expression. In line with the significant results seen in HeLa-Ohio cells, hsa-miR-140-3p was associated with multiple genes (STAG2 and SMARCA2) involved in the immune response, suggesting this miRNA is strongly associated with antiviral responses and viral progression. This initially suggests that hsa-miR-140-3p could represent a strong potential biomarker for RV16 infection. SMARCA2, while having a relatively low VarElect GeneCards score, has been shown to support antiviral responses, including activating IFN-stimulated genes (ISGs) and being involved in the intrinsic apoptotic pathway. Deficiency of SMARCA2 has been shown to activate signalling pathways such as cGAS-STING (Cyclic-GMP-AMP Synthase-Stimulator of Interferon Genes) and JAK-STAT (Janus kinase/signal transducers and activators of transcription), supporting the hosts antiviral responses (Dornfeld *et al*. 2018; Dudek *et al*. 2018). STAG2 has also been shown to be regulated by hsa-miR-140-3p through the mirDIP database and regulates ISG expression. This further supports the suggestion that hsa-miR-140-3p has complex roles during RV16 infection and could be a useful biomarker at multiple time points.

Downregulation of hsa-miR-122-5p is also critical and represented here via CD40LG. CD40LG, encodes a protein expressed on T cells and regulates B cell function (Aytekin and Vaqar 2023). Due to this, reduced CD40LG would weaken the adaptive antiviral response and allow viral replication, suggesting that RV16 potentially downregulates hsa-miR-122-5p to supress adaptive immune cell function.

Finally, hsa-miR-155-5p represents a strong candidate biomarker. From the program and literature search, we found that OLFML3 was strongly correlated to RV16 infection. OLFML3 has been linked to RV14 infection, being involved in IFN-I responses in a SOCS3 (suppressor of cytokine signalling 3) dependent manner, suggesting RV targets hsa-miR-155-5p to reduce OLFML3 and ultimately reduce host immune responses (Mei *et al*. 2021). This supports a role of hsa-miR-155-5p in RV infection, offering potential as a specific biomarker helping to determine the stage of infection.

Overall, these results suggest that RV16 targets specific miRNAs to promote its infection, and this could be mediated via specific gene expression patterns aimed at dampening down the immune response.

## Discussion

Overall, these results indicated that, during RV16 infection, there is differential expression of miRNAs at varying time points. Assessing a large range of early and late time points across 3 miRNAs and U6 snRNA allowed for a broad analysis and identification of potential biomarkers for RV16 infection. Interestingly, of these, hsa-miR-140-3p and U6 were found to show the most significance, in contrast to studies in RV1B where hsa-miR-155-5p was found to be a strong marker of infection severity and progression (Bondanese *et al*. 2014). This suggests significant variations in how major and minor RV virus’ may modify the miRNA and gene landscape but shows the value of these miRNAs as diagnostic biomarkers for RV infection. It’s likely there will be some strain specificities as well as overlaps meaning there would be a panel that can be used universally with updates easily added into any diagnostic test to pull out specific differences. Combined with traditional PCR based tests this could be very attractive for the future prevention of RV infections.

When assessing miRNAs as biomarkers for RV infection, it is important to understand what makes an effective biomarker, and the roles their gene targets play in viral infection and the host immune response. Overall, despite cell specific differences, there are some commonalities making 3 of these miRNAs attractive. hsa-miR-155-5p presents an attractive biomarker for infection, being strongly downregulated in the supernatant during the peak of viral replication (15-hours). hsa-miR-155-5p has been shown to influence a wide array of pathways across viral infection and the immune response, being essential for antiviral Th1 and Th2 responses which can limit inflammation and viral replication. Out of all genes assessed here, OLFML3 is the most directly relevant to RV infection, being shown to antagonise IFN-I in a SOCS3 dependent manner. Knockout of this gene has been shown to reduce RV14 infection in H1-HeLa cells, suggesting RV directly downregulates hsa-miR-155-5p, increasing OLFML3 expression, and ultimately blocking IFN-I responses (Mei *et al*. 2021). A recent study supported this, finding hsa-miR-155-5p overexpression to be restrictive to RV-1B infection (Bondanese *et al*. 2014), and is consequently overexpressed in RV-infected airways. Our results support the diagnostic benefit of hsa-miR-155-5p as expression levels remained relatively stable until 15-hours post-infection, with RV16 likely targeting it to drive OLFML3 expression, preventing early antiviral responses during peak replication. The results in other studies further support this (Arroyo *et al*. 2020), and along with our

work suggest this could represent a promising diagnostic biomarker of RV16 infection at crucial replication time points. In Hep2 cells, a similar effect is seen, however there is clear upregulation at 24-hours post-infection, far larger than that seen in HeLa-Ohio cells, suggesting there may be slight differences in infection dynamics between the 2 cell types. However, there seems to be a more consistent downregulation suggesting that there is a wider disruption of the antiviral response. However, the results still support our findings and further suggest that hsa-miR-155-5p could be a promising diagnostic biomarker.

hsa-miR-140-3p showed differences in expression between the 2 cell lines but consistency at 15-hours. hsa-miR-140-3p has been strongly associated with regulation of gene expression, involvement in apoptosis, and promoting viral replication. Looking first at SMARCA2, part of the BAF (Barrier-to-autointegration factor) chromatin remodelling complex, this gene has been shown to be directly related to supporting antiviral responses, including activating ISGs (Dornfeld *et al*. 2018; Dudek *et al*. 2018). In late-stage infection (24-hours), C-terminally cleaved SMARCA2 accumulates. This cleavage removes domains important for nuclear localisation and chromatin binding, meaning SMARCA2 cannot accumulate in the nucleus and fully activate ISG expression. SMARCA2 is also targeted by cellular caspases downstream of the intrinsic apoptotic pathway, further underlining its role in cell death pathways and restricting viral replication (Dudek *et al*. 2018). Despite cell specific differences, the significant downregulation at 15-hours in both cell types was key. This downregulation likely reflects the viruses attempt to target genes such as SMARCA2 to promote viral replication. We can suggest that RV16 causes the C-terminal cleavage of SMARCA2 and leads to its cytoplasmic accumulation, weakening the IFN response and coinciding with the changes seen for hsa-miR-155-5p (Dudek *et al*. 2018). However, this is likely transient and requires other genes to be activated as it could be overcome.

In addition to SMARCA2, STAG2 has also been shown to be regulated by hsa-miR-140-3p. STAG2 is an integral part of the cohesion complex, regulating separation of sister chromatids during cell division, and having strong links to cancer progression. As well as this, STAG2 has been observed to play a central role in viral infection including rotavirus and porcine deltacoronavirus. In rotavirus, STAG2 deficiency induces IFN responses via the cGAS-STING pathway, ultimately restricting viral infection. This subsequently activates the JAK-STAT signalling pathway and ISG expression (Ding *et al*. 2018). This same effect is observed in porcine deltacoronavirus (Wu *et al*. 2022). When assessing this in relation to RV16 and the above results, it can be suggested that the virus attempts to upregulate STAG2 to prevent an effective antiviral response being mounted, allowing establishment of viral infection. In contrast to SMARCA2, it is likely that hsa-miR-140-3p downregulates STAG2, meaning it would be beneficial for RV16 to decrease expression of this miRNA to prevent ISG and IFN expression, ultimately allowing it to effectively replicate and progress. At 15-hours, this effect is seen, where hsa-miR-140-3p is significantly downregulated compared to 2-hours, upregulating STAG2, preventing IFN responses and ISG expression. However, as the infection progresses, hsa-miR-140-3p levels begin to recover, which is in line with a more robust immune response starting. While no studies appear to draw a clear link between hsa-miR-140-3p and STAG2 expression, especially in the context of viral infection, it is likely that there is a direct correlation between these which could be a target of future research.

From this, it can be seen that STAG2 and SMARCA2, while both being regulated by the same miRNA, have directly contrasting roles with one being antiviral (SMARCA2) and the other supporting viral infection (STAG2). Therefore, it is no surprise that the virus targets these genes, as seen through the significant downregulation of hsa-miR-140-3p at 15-hours post-infection, as RV16 attempts to dampen the host response to support replication and progression. Despite more studies being needed our results do also suggest that hsa-miR-140-3p could also represent a potential diagnostic biomarker for RV16 and, combined with hsa-miR-155-5p, could be effective within a biomarker panel.

Overall, from the above analysis and results, hsa-miR-140-3p and hsa-miR-155-5p present as the most reliable and specific biomarkers for infection, based on their gene targets and overall role in viral infections and the immune response throughout infection. While hsa-miR-122-5p does have some role within viral infection, and regulates CD40LG, in both cell lines, its role is not as well established as other miRNAs. However, some specificities can be drawn. CD40LG encodes a protein that is expressed on T cells and regulates B cell and dendritic cell (DC) function with mutations promoting higher susceptibility to infection (Aytekin and Vaqar 2023). Consequently, increased CD40LG expression is strongly linked with severe viral infection, via enhanced DC activation and, in the context of viruses, CD8+ T cell activation. For example, in SARS-CoV-2, higher CD40LG was observed in intensive care patients and detectable in plasma samples (Blot *et al*. 2020). In relation to our results, the initial upregulation of hsa-miR-122-5p reflects these processes and the initial response upon recognition of the virus by the host. At this early stage of viral entry, it is likely hsa-miR-122-5p upregulation is an attempt to promote DC activation and hence an appropriate T cell response via CD40LG expression. This then gradually declines until 15-hours potentially due to virus induced processes, leading to reduced viral clearance. Due to this, it is likely the observed changes in this miRNA’s expression are most likely due to the host attempting to release hsa-miR-122-5p to activate DCs, as well as priming CD8+ T cells, enhancing viral responses (Moffett *et al*. 2017). In healthy patients, this response may be robust and successful at either clearing or restricting RV16 infection, however in patients with asthma or COPD there may be a weaker response with less CD8+ T cell recruitment, preventing viral clearance. This supports hsa-miR-122-5p as having a key role in RV infection, but more restrictive to a host directed response with limited and indirect input from the virus. Furthermore, unlike our other miRNAs it doesn’t show a broad degree of modification that is consistent across cell types restricting its use as a diagnostic biomarker. Despite this, it is important not to rule out hsa-miR-122-5p as a biomarker or therapeutic target, as its early 1.5-fold upregulation, and downregulation at 15-hours post-infection suggests that RV16 does target this miRNA. As with hsa-miR-155-5p, it may be more useful within a panel of related biomarkers than alone, but clearly further work is needed here.

Despite these results our work does have some limitations. Firstly, this project primarily focused on only a small number of miRNAs, meaning further miRNAs need to be assessed. In addition, only cell lines were used, meaning there may be differences between the miRNA changes seen here, and that seen in patient cells. Future research will use primary cells and patient samples. Finally, the analysis, while providing insightful information into gene targets and miRNA pathways, is not mechanistic but this will be the basis of the next studies.

Overall, the results here suggest that all three miRNAs examined play a role in RV16 infection that could occur through regulation of gene expression involved in viral replication and signalling pathways such as cGAS-STING and JAK-STAT. We can suggest from our analysis that miR-155 and 140 represent the strongest potential biomarkers that can be taken forwards in further studies in the future. With future work assessing mechanisms and specificities, these miRNAs could be used as biomarkers either independently or as part of a panel, for early identification of RV16 infection.

## Acknowledgements

We are grateful to Pax Bosner for providing initial conceptual modelling scripts on which this preprint developed. We would also like to thank Genecards for allowing us to use their API.

